# Heat Tolerance is Affected by the Gut Microbiota in a Vertebrate Ectotherm

**DOI:** 10.1101/2023.09.07.556683

**Authors:** Jason W. Dallas, Anna Kazarina, Sonny T. M. Lee, Robin W. Warne

**Affiliations:** Southern Illinois University, School of Biological Sciences, 1125 Lincoln Dr., Carbondale, IL 62901-6501; Kansas State University, Division of Biology, 1717 Claflin Rd, Manhattan, KS 66506

**Keywords:** CT_max_, Thermal Acclimation, Gut microbiota, Wood frogs, Microbiome manipulation

## Abstract

The gut microbiota is known to influence and have regulatory effects in diverse physiological functions of host animals, but only recently has the relationship between host thermal biology and gut microbiota been explored. Here, we examined how early-life manipulations of the gut microbiota in larval amphibians influenced their critical thermal maximum (CT_max_) at different acclimation temperatures. We removed the resident microbiome on the outside of wild-caught wood frog (*Lithobates sylvaticus*) egg masses via an antibiotic wash, and then either maintained eggs without a microbiota or inoculated eggs with pond water or the intestinal microbiota of another species, green frogs (*L. clamitans*), that have a wider thermal tolerance. We predicted that this cross-species transplant would improve the CT_max_ of the recipient wood frog larvae relative to the other treatments. In line with this prediction, green frog-recipient larvae had the highest CT_max_ while those with no inoculum had the lowest CT_max_. Both the microbiome treatment and acclimation temperature significantly influenced the larval gut microbiota communities and alpha diversity indices. Green frog inoculated larvae were enriched in Rikenellaceae relative to the other treatments, which produce short-chain fatty acids and could contribute to greater energy availability and enhanced heat tolerance. Larvae that received no inoculation had higher relative abundances of potentially pathogenic *Aeromonas* spp., which negatively affects host health and performance. Our results are the first to show that cross-species gut microbiota transplants alter heat tolerance in a predictive manner. This finding has repercussions for the conservation of species that are threatened by climate change and demonstrates a need to further explore the mechanisms by which the gut microbiota modulates host thermal tolerance.

## Introduction

The host microbiota represents a complex biological community consisting of bacteria, fungi, and viruses that has received increased focus over recent decades as its highly influential role on host physiology has become elucidated (Kohl and Carey, 2016; McFall-Ngai et al., 2013; Warne and Dallas, 2022). Resultantly, how the host microbiota responds to changes in environmental variables that are intimately tied to organismal biology, such as temperature, are of great interest. There is correlative evidence that environmental temperatures alters both the gut microbiota compositional and functional diversity in vertebrates and invertebrates (Hassenrück et al., 2021; Jaramillo and Castaneda, 2021; Kohl and Yahn, 2016; Li et al., 2020; Moghadam et al., 2018; Onyango et al., 2020; Wang et al., 2018; Woodhams et al., 2020). In their review, Sepulveda and Moeller (2020) found that warming temperatures are associated with increased relative abundance of Proteobacteria in invertebrates and a reduction in Firmicutes in vertebrates, and these temperature-dependent changes in the microbiota community affect varying aspects of host fitness and performance (Fontaine et al., 2018; Horváthová et al., 2019; Kikuchi et al., 2016; Prado et al., 2010). Therefore, the risks of climate change, which are expected to worsen over the coming decades (IPCC, 2021), extend to the host microbiota and represents an important topic of study.

A fundamental aspect of an ectotherm’s biology is their thermal tolerance, as it limits the temporal and spatial range they can occupy (Adolph and Porter, 1993; Hoffmann et al., 2013; Huey, 1982). Thermal tolerance of ectotherms is typically estimated via critical thermal (CT) limits that denote the endpoints of their thermal performance curve (TPC) where organismal performance reaches zero (Huey and Stevenson, 1979). Species with higher CT maxima (CT_max_) are expected to be less susceptible to warmer environments enabling them to persist in the projected climates (Gilbert and Miles, 2017; Huey et al., 2012; Roeder et al., 2021). Therefore, we suggest that a warmer CT_max_ is beneficial to many populations going forward. The factors that underlie organismal CT_max_, and to a broader extent, heat death in ectotherms remains debated. Several proposed mechanisms include perturbations in the plasma membrane (Bowler, 2018), a decline in aerobic scope (Pörtner, 2002), loss of cardiovascular function (Somero, 2010), and disruption in mitochondrial activity (Chung and Schulte, 2020). Due to the role of CT_max_ in promoting species persistence, there is foremost importance to exploring the different aspects of an individual’s environment and biology that can elevate CT_max_.

One potential, yet relatively understudied, factor of organismal biology that can influence host heat tolerance is the host microbiome. Across several invertebrate and vertebrate species, there are correlative links between the host microbiome and changes in CT_max_ (Baldassarre et al., 2022; Doering et al., 2021; Fontaine et al., 2022; Moeller et al., 2020). For example, an increasing prevalence of the *Anaerotignum* genus positively improved the CT_max_ of western fence lizards *Sceloporus occidentalis* (Moeller et al., 2020). Additionally, Doering et al. (2021) showed that transplanting the microbiome of coral populations from thermally fluctuating waters to conspecifics from thermally stable environments enhanced their bleaching resistance, indicative of greater heat tolerance. There are several potential pathways through which the host microbiota can elevate host CT_max_. For instance, mitochondrial enzyme activity was reduced in larval anurans with an experimentally-depleted microbiota (Fontaine et al., 2022), which can decrease aerobic scope at high temperatures and limit CT_max_ (Pörtner, 2002). Furthermore, warm-acclimated microbiota community exhibit higher expression of reactive oxygen scavengers (Fontaine and Kohl, 2023; Ziegler et al., 2017) that can limit the damage reactive oxygen species impart on mitochondrial mechanisms at high temperatures (Christen et al., 2018; Sokolova, 2023). Beyond enhanced CT_max_, members of the host microbiota can promote enhanced host fitness in warmer temperatures as demonstrated in pea aphids *Acyrthosiphon pisum* hosting an essential bacterial symbiont (*Buchnera aphidicola*) (Montllor et al., 2002; Zhang et al., 2019). These examples suggest that the host microbiome confers benefits to heat tolerance in the host that could facilitate survival in warmer environments.

Amphibians are imperiled by climate change as increases in temperature and unpredictable shifts in precipitation will restrict activity periods and place species at risk of experiencing heat stress (Campbell Grant et al., 2020; Greenberg and Palen, 2021; Hoffmann et al., 2021; Sunday et al., 2014). Based on the risks associated with warming temperatures, we sought to examine how early-life manipulation of the gut microbiota in a geographically widespread anuran influenced its heat tolerance. Specifically, we used a cross-species microbiota transplant to demonstrate that species-specific differences in thermal tolerance is mediated, at least partially, by the gut microbiota. While cross-species microbiota transplants have not been examined in terms of heat tolerance, Warne et al. (2019) showed that egg masses of wood frogs (*Lithobates sylvaticus*) inoculated with the gut microbiota of larval bullfrogs (*L. catesbeianus*) displayed higher growth and developmental rates as well as enhanced disease resistance compared to those inoculated with a conventional wood frog microbiota.

To evaluate if gut microbiota and inter-specific transplants influence thermal tolerance, we stripped the resident microbiota of field-collected wood egg masses using antibiotics and separated these “sterilized” eggs into three microbiota treatments. Following the antibiotic wash, eggs were either inoculated with pond water from the collection site, the gut microbiota of field-collected larval green frogs (*L. clamitans*), or received no inoculation. As green frogs breed in warmer temperatures than wood frogs and their larvae are active over the summer months while wood frogs metamorphose prior to the summer (Hulse et al., 2001), green frog larvae have been shown to exhibit a CT_max_ of more than 1.5°C higher than wood frog larvae (Katzenberger et al., 2021). Resultantly, we predicted that inoculating wood frog eggs with the gut microbiota of larval green frogs would increase the heat tolerance of the recipient. We also evaluated if a short-term acclimation period would alter the gut microbiota. Most studies measuring the effects of different acclimation temperatures on the gut microbiota exceed 10 days (e.g., Fontaine et al., 2022; Moeller et al., 2020; Zhu et al., 2021), and the impact of a short-term thermal acclimation period has not been thoroughly explored. Therefore, we sought to identify how the gut microbiota changes in response to a brief acclimation period of three days and if there was an interactive effect between acclimation temperature and microbiota treatment on wood frog CT_max_.

## Materials and Methods

### Field Collection

Four freshly laid (< 36 hours old) wood frog egg masses were collected from wetlands in Jackson Co., IL, under an Illinois Department of Natural Resources permit (HSCP 19-03). Additionally, 10 larval green frogs were collected from Southern Illinois University Carbondale research ponds in Jackson Co., IL. All experimental procedures were approved by the Southern Illinois University Institutional Animal Care and Use Committee (22–008).

### Gut microbiome treatments

On the day of collection, eggs were rinsed in antibiotics following a previously published protocol (Warne et al., 2017; Warne et al., 2019). Briefly, the egg masses were separated into sterile 50 mL tubes (∼ 20 eggs/tube), rinsed three times with 40mL of autoclaved, aerated carbon-filtered water. The eggs were then sterilized by exposure to 500 µL penicillin-streptomycin (10,000 U/mL; Life Technologies #15140-122), 200 µL of kanamycin sulphate (25 µg/mL; Life Technologies #11815-032) and 50 µL of amphotericin B solution (250 µg/mL; Sigma-Aldrich #A2942) for 4 hours on a nutator. The sterilized eggs were then triple rinsed with sterile water and placed into sterilized 6 L plastic containers containing 3 L of carbon-filtered sterile water and aerated with a HEPA inline filter disc (0.3 µm pore, Whatman Inc.) overnight. Additionally, we retained eggs from all four masses that were not exposed to any antibiotics.

The following day, the antibiotic-rinsed eggs were collected and separated into three different treatments: 1) wood frog (WF), 2) no-inoculum (NI), and 3) green frog (GF). The WF treatment eggs were placed in sterile 50 mL tubes and received 40 mL of pond water that was previously mixed with the unmanipulated eggs with the intention of re-inoculating them with their natural microbiota. The NI treatment received no further manipulation, and all microbial colonization of this group was assumed to be environmentally acquired post-hatching. For the GF treatment, intestinal tracts were dissected from the larval green frogs (Gosner stage 27–30; Gosner, 1960), homogenized in 40 μL of autoclaved water, and added to sterile 50 mL tubes with the sterilized wood frog eggs. All treatments were mixed with their respective inoculum for 30 minutes on a nutator and were then returned to their respective containers (N = 16 per treatment).

Larvae hatched within four days of the microbiota treatments and were allowed to feed on the inoculated egg jelly for two days. Bubblers were removed one week after hatching. Initial feedings consisted of autoclaved algal flakes (Bug Bites Spirulina Flakes, Fluval Aquatics, Mansfield, MA, USA) after which they were fed crushed alfalfa pellets twice weekly. Water was changed weekly with aerated, carbon-filtered water.

### Critical Thermal Maximum Assay

Over a four-day period, we randomly collected 40 larvae from each microbiota treatment. All larvae were staged, weighed, and transferred to individual 750 mL plastic containers filled with 600 mL of aged (>24 hours) aerated, carbon-filtered water. To reduce the effect of individual containers, only one to three larvae were collected from each container per day. Across all individuals, larval stages ranged from 27 – 36, and there was no difference among the treatments in terms of stage (F_2,107_ = 0.92, P = 0.40) or log-transformed mass (F_2,107_ = 0.75, P = 0.48), but collection day influenced larval stage (F_3,107_ = 4.44, P = 0.006) but not mass (F_3,107_ = 1.96, P = 0.13) and there was no interaction. Lastly, larvae (n = 20 per microbiota treatment) were then split into two acclimation temperatures, low (15°C ± 0.2) and high (23°C ± 0.3), for a three-day acclimation period during which individuals were fasted. Following an attempt to measure the CT minimum of larvae from the low acclimation temperature, the sample size of was reduced to N = 18.

After the acclimation period, larvae were staged and weighed then placed in individual 125 mL flasks filled with 75 mL of aged, aerated, carbon-filtered water. Larvae were then submerged in a hot water bath (Isotemp 220, Fischer Scientific) and given 5 minutes to acclimate prior to beginning the assay. In each bath, there were eight flasks containing two individuals from each treatment. To record CT_max_, temperatures increased ∼0.7°C per minute from a starting temperature (mean ± 1 standard error) of 19.4 ± 0.2°C. Beginning at 33°C, larvae were prodded with a spatula every 30 seconds until they failed to respond to the stimulus. At this point, a thermocouple probe (Physitemp BAT-12) was placed adjacent to the larvae and the water temperature was recorded, which represented larval CT_max_. Flasks were then placed in a bath of room temperature water to facilitate larval recovery. While most larvae recovered within 3 minutes, 6 larvae died from the heat shock and there was no treatment effect (χ^2^ = 3.31, df = 2, P = 0.19), and two individuals died prior to the assay; all were removed from further analyses.

To test for acclimation effects on gut microbiomes, four larvae were selected for microbial sequencing from each of the six groups (three microbiota treatments x two acclimation temperatures) after three days of temperature acclimation. We did not measure the CT_max_ of these 24 larvae to eliminate potential confounding effects of heat shock on the microbiota community. All larvae were euthanized via snap-freezing in -80°C ethanol and stored at -80°C until intestinal extraction.

### Gut Microbiota DNA Extraction

The entire intestinal tract was removed and placed into an autoclaved 1.5 mL microcentrifuge tube. To reduce cross-contamination, forceps were rinsed in 70% ethanol and then flame-sterilized prior to intestine removal. Additionally, intestines were briefly rinsed with autoclaved water to remove any transient bacteria. To improve DNA extraction, the intestines were homogenized using sterilized forceps and then stored at -80°C. Microbial DNA was extracted using the GenCatch™ Plasmid DNA Mini-Prep Kit (Epoch Life Science) following manufacturer instructions with minor modifications: The volume of Proteinase K was increased to 25 μL and the initial incubation at 60°C was increased to 3 h to improve cellular digestion. Lab controls were used to identify any microbial DNA present in the extraction kit reagents. All extracted DNA samples were stored at -80°C prior to sequencing.

### Gut Microbiota Analyses

We used QIIME 2 v. 2021.4 (Bolyen et al., 2019) to process a total of 7,573,956 raw sequences, resulting in 4,293,502 bacterial sequences after quality control for 87 samples. Note that only 24 of these 87 samples are for this current study, while the remaining samples are a parallel study of larval wood frogs. We used QIIME 2 plugin cutadapt (Martin, 2011) to remove the primer sequences. Any sequences with ambiguous bases, with no primer, with greater than 0.1 error rate mismatch with primer or any mismatches to the sample-specific 12 bp molecular identifiers (MIDs) were discarded. Following initial quality control, we used DADA2 (Callahan et al., 2016) to truncate the reads to length where the 25th percentile of reads had a quality score below 15 (Forward 271 and Reverse 277). We used the pre-trained SILVA database (v. 138) in QIIME 2 for taxonomic assignment of the bacteria. Sequences were annotated to amplicon sequence variants (ASVs), and any unknown or unclassified ASVs were removed from downstream analysis. We rarefied the data set to 10,000 reads per sample (resulting in 870,000 high quality sequences variants in the 87 samples) to minimize biases resulting from differences in sequencing depth among samples before estimating diversity indices and downstream analyses (Gihring et al., 2012).

### Statistical Analyses

All statistical analyses were conducted using R version 4.1 (Team, 2021). Larval heat tolerance was assessed via a linear mixed model using the lmer package (Kuznetsova et al., 2017) with CT_max_ as the response variable and both microbiota treatment and acclimation temperature as fixed effects. We included larval mass, which was square root transformed to improve normality, as a covariate and the location of the 125 mL flask in the hot water bath as a random effect to account for potential differences in heating rates. In the results, CT_max_ is reported as mean ± one standard error along with sample size.

Gut microbiota alpha and beta diversity metrics were analyzed using the vegan package (Oksanen et al., 2022). We compared three alpha diversity metrics (ASV richness, Shannon index, and Simpson index) using two-way ANOVAs with microbiota treatment, acclimation temperature, and their interaction as predictive variables and Gosner stage as a covariate. We used the adonis2 function to perform PERMANOVAs, with 999 permutations, using Bray-Curtis and Jaccard distance matrices based on microbiota treatment, acclimation temperature, and their interaction along with Gosner stage as a covariate. Using the same distance indices, we compared intragroup heterogeneity via permutation tests with 999 permutations and the betadispr function.

We determined significant differences in bacterial phyla and families across the microbiota treatments and acclimation temperatures using the MaAsLin2 package (Mallick et al., 2020). We used an arcsine-square root transformation to improve normality of the relative abundance data. To reduce the risks of false positives, we corrected P-values using the Benjamini-Hochberg false discovery rate method.

## Results

### Heat tolerance differences between gut microbiota treatments

We found that larval wood frog CT_max_ was affected by gut microbiota treatment and acclimation temperature, although the differences in heat tolerance were small (Fig. 1). In line with our prediction, microbial treatment, controlling for body mass and acclimation temperature, influenced CT_max_ (Table 1; GLMM, χ^2^ = 14.86, P < 0.001). GF larvae had the highest CT_max_ (38.3°C ± 0.2, N = 13) while WF and NI larvae (37.7°C ± 0.1, N = 12) had the lowest per post-hoc pairwise comparisons within the low acclimation temperature (P = 0.051). Acclimation temperature had a pronounced effect on CT_max_ in the expected direction, when controlling for body mass and microbiota treatment, with those acclimated to warmer temperatures exhibiting higher heat tolerance (Table 1; GLMM, χ^2^ = 19.98, P < 0.0001). A post-hoc analysis found that only NI larvae showed a significant increase (P = 0.037) in CT_max_ between the low and high acclimation temperatures of 0.6°C, while the GF (0.3°C) and WF (0.4°C) treatments had smaller increases. There was no interaction between gut microbiota treatment and acclimation temperature, and the differences in CT_max_ were independent of body mass (Table 1).

**Figure 1:**
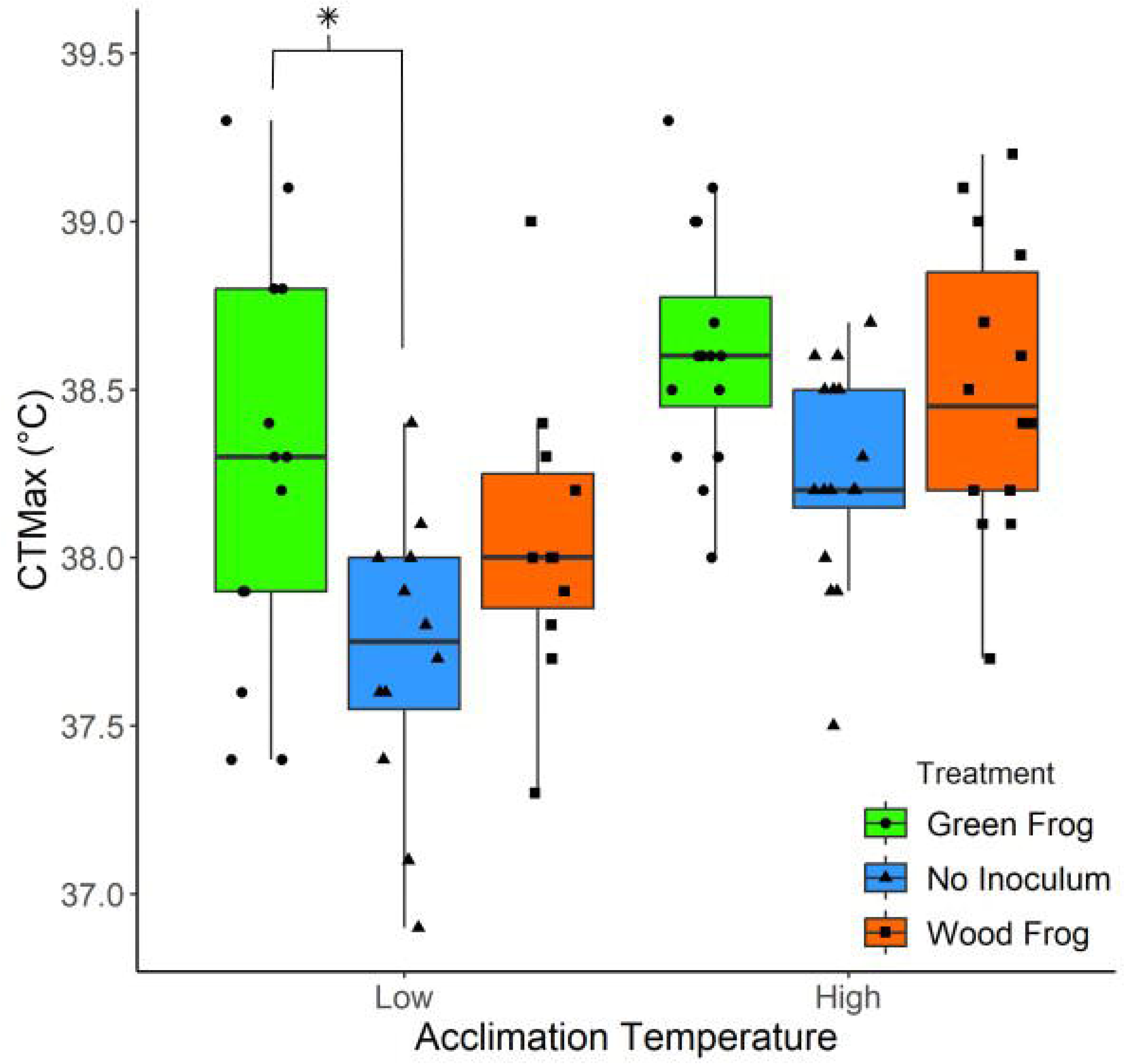
The critical thermal maximum (CT_max_) of larval wood frogs was dependent on their gut microbiota treatment and their acclimation temperature. Larvae acclimated to a higher temperature had a higher CT_max_ independent of microbiota treatment and the green frog larval treatment had higher CT_max_. The asterisk represents a near-significant pairwise difference (P = 0.51) within the acclimation temperature. Center lines within boxplots represent the median and the boxes denote the interquartile range with whiskers representing 1.5× the upper or lower quartile.

**Table 1:** Effects of square root transformed body mass, microbiota treatment, acclimation temperature, and their interactions on larval wood frog critical thermal maximum from a generalized linear model.

| Variable | $\chi^2$ | df | *P* |
| --- | --- | --- | --- |
| √Mass | 0.16 | 1 | 0.69 |
| Microbiota Treatment | 14.86 | 2 | **< 0.001** |
| Acclimation Temperature | 19.98 | 1 | **< 0.0001** |
| √Mass x Microbiota Treatment | 1.04 | 2 | 0.6 |
| √Mass x Acclimation Temperature | 0.0032 | 1 | 0.96 |
| Microbiota Treatment x Acclimation<br>Temperature | 0.94 | 2 | 0.62 |
| √Mass x Microbiota Treatment x<br>Acclimation Temperature | 0.92 | 2 | 0.66 |

### Alpha Diversity

We analyzed a total of 1,221,946 sequences that were assigned to 145 unique ASVs across the 24 intestinal samples. Across the three gut microbiota treatments, ASV richness was lowest in the WF group and highest in the GF group, but there was no significant treatment or acclimation temperature effect (Fig. 2A). Using Shannon’s and Simpson’s indices, alpha diversity was significantly different across microbiota treatments, acclimation temperature, and larval stage. The WF larvae had significantly lower diversity metrics than both GF and NI larvae, and those acclimated to the high temperature had higher diversity, although the difference was minor (Figs. 2B and 2C). Across both indices, diversity increased with larval stage, but Shannon’s index model had a near-significant interaction of stage and treatment as NI larvae showed no change in diversity across different stages (Fig. 2B).

**Figure 2:**
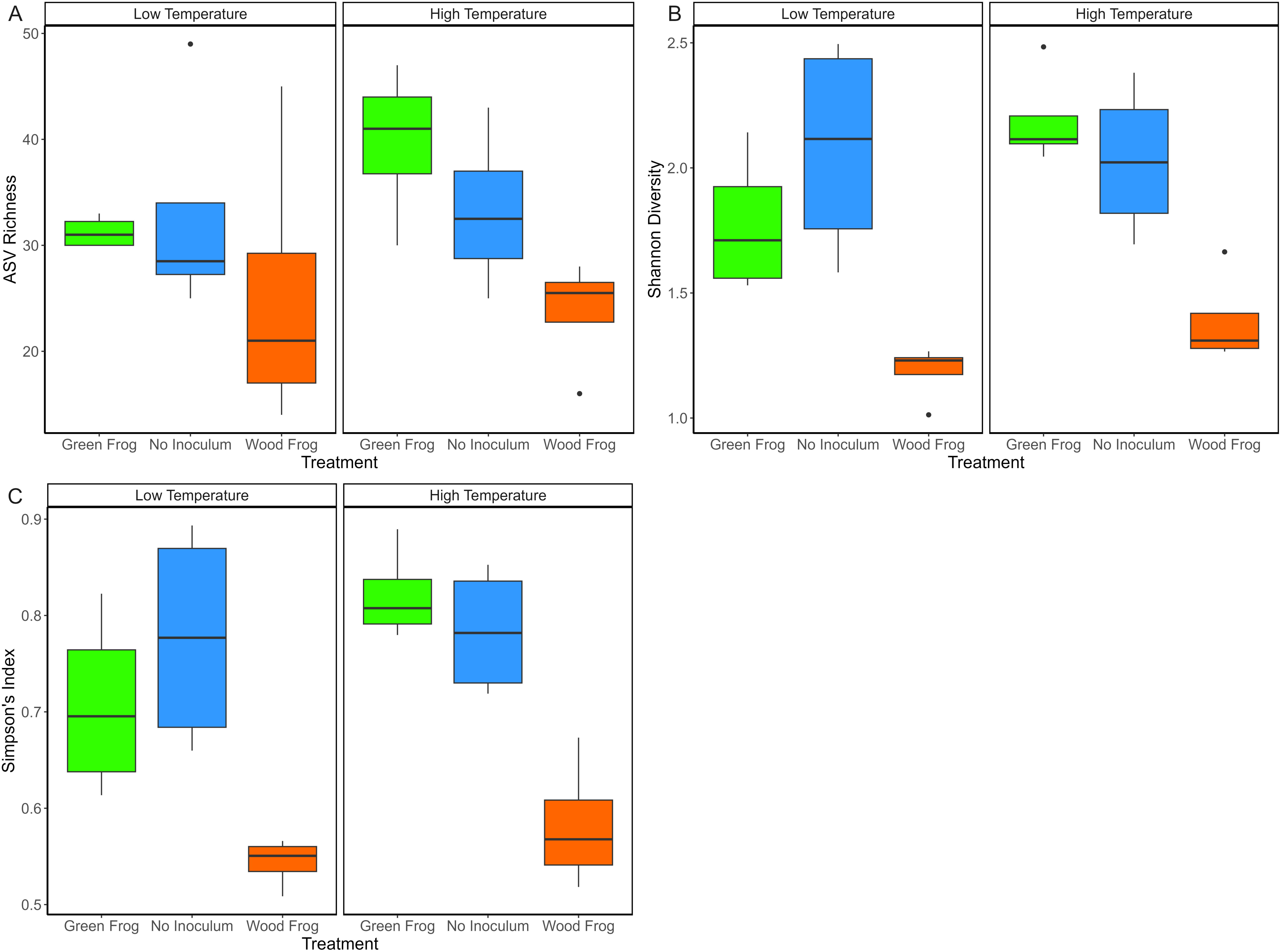
The effect of gut microbiota treatments and acclimation temperature on alpha diversity indices of larval wood frogs: A) ASV Richness, B) Shannon Index, and C) Simpson Index. For both Shannon and Simpson diversity, the wood frog treatment had significantly lower values than the other treatments at both acclimation temperatures (Tukey HSD P < 0.05), but approached significance (P < 0.08) when compared against the low temperature green frog treatment. Center lines within boxplots represent the median and the boxes denote the interquartile range with whiskers representing 1.5× the upper or lower quartile.

### Beta Diversity

The community composition and membership of the wood frog gut microbiota was varied with respect to microbiota treatment, acclimation temperature, and larval stage (Fig. 3). Adonis PERMANOVAs identified significant effects of the microbiota treatments (Bray-Curtis and Jaccard distances, respectively: F_2,14_ = 4.22, P < 0.001; F_2,14_ = 2.98, P < 0.001), temperature (F_1,14_ = 4.41, P < 0.001; F_1,14_ = 3.27, P < 0.001), and Gosner stage (F_1,14_ = 2.90, P = 0.009; F_1,14_ = 2.09, P = 0.013). For both distance methods, there were no significant interactions. The PCoA plots of Bray-Curtis and Jaccard distances showed moderate overlap among the microbiota treatments and acclimation temperatures (Fig. 3). While the NI larvae at the low acclimation temperature was the most variable, within-group dispersion was similar for both Bray-Curtis (F_5,18_ = 1.15, P = 0.36) and Jaccard (F_5,18_ = 1.11, P = 0.41) distance analyses. Among the most prominent phyla, variation along PC1 of both distance methods was negatively related to Bacteroidota abundance while Firmicutes and Proteobacteria abundances had positive effects. Actinobacteria accounted for a majority of variation along PC2 being negatively related in the Bray-Curtis distance PCoA and positively related in the Jaccard distance PCoA.

**Figure 3:**
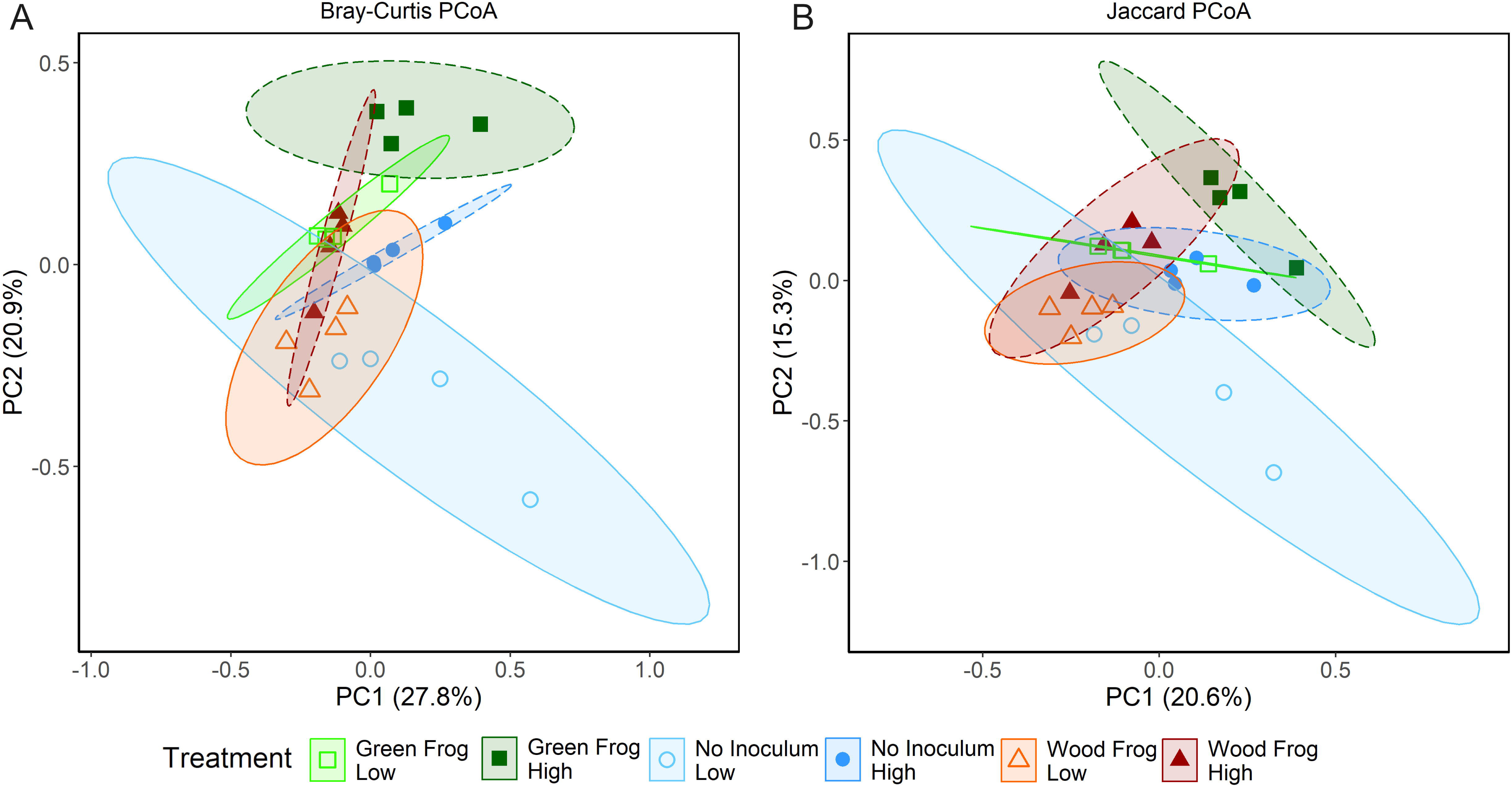
Principal coordinate analyses (PCoA) of the gut microbiota community structure (A) and membership (B) of larval wood frogs that underwent early life microbiota treatments and were acclimated to either low or high temperatures. Larvae from the low acclimation temperature are indicated by hollow symbols and lighter colors relative to high temperature acclimated larvae. PCoAs were based on all ASVs. Percentages on axes titles represent the variance explained by the eigenvector. Each point represents the gut microbiota of an individual wood frog.

Across all wood frog larvae, gut bacterial communities were dominated by three phyla: Bacteroidota, Firmicutes, and Proteobacteria with relatively lower abundances of Actinobacteria, Desulfobacterota, and Verrucomicrobiota (Fig. 4). Across both acclimation temperatures, the relative abundance of Bacteroidota was lower in NI larvae (mean ± 1 SE: 38.7 ± 6.6%) compared to the GF (53.4 ± 5.7%) and WF larvae (66.0 ± 2.4%), but a higher relative abundance of Proteobacteria (38.7 ± 6.1%) with respect to GF (22.1 ± 3.8%) and WF larvae (16.8 ± 2.8%). However, these differences were not statistically significant (MaAsLin2, across all treatments corrected P > 0.09; Table S1). The relative abundance of these major phyla was unaffected by the acclimation temperature indicating their resistance to the short-term acclimation period (MaAsLin2, corrected P > 0.72). Across all treatments, Actinobacteria was enriched under the low acclimation treatment (1.1 ± 0.3%) relative to the high temperature (0.45 ± 0.2%), although the difference only approached statistical significance (MaAsLin2, corrected P = 0.067).

**Figure 4:**
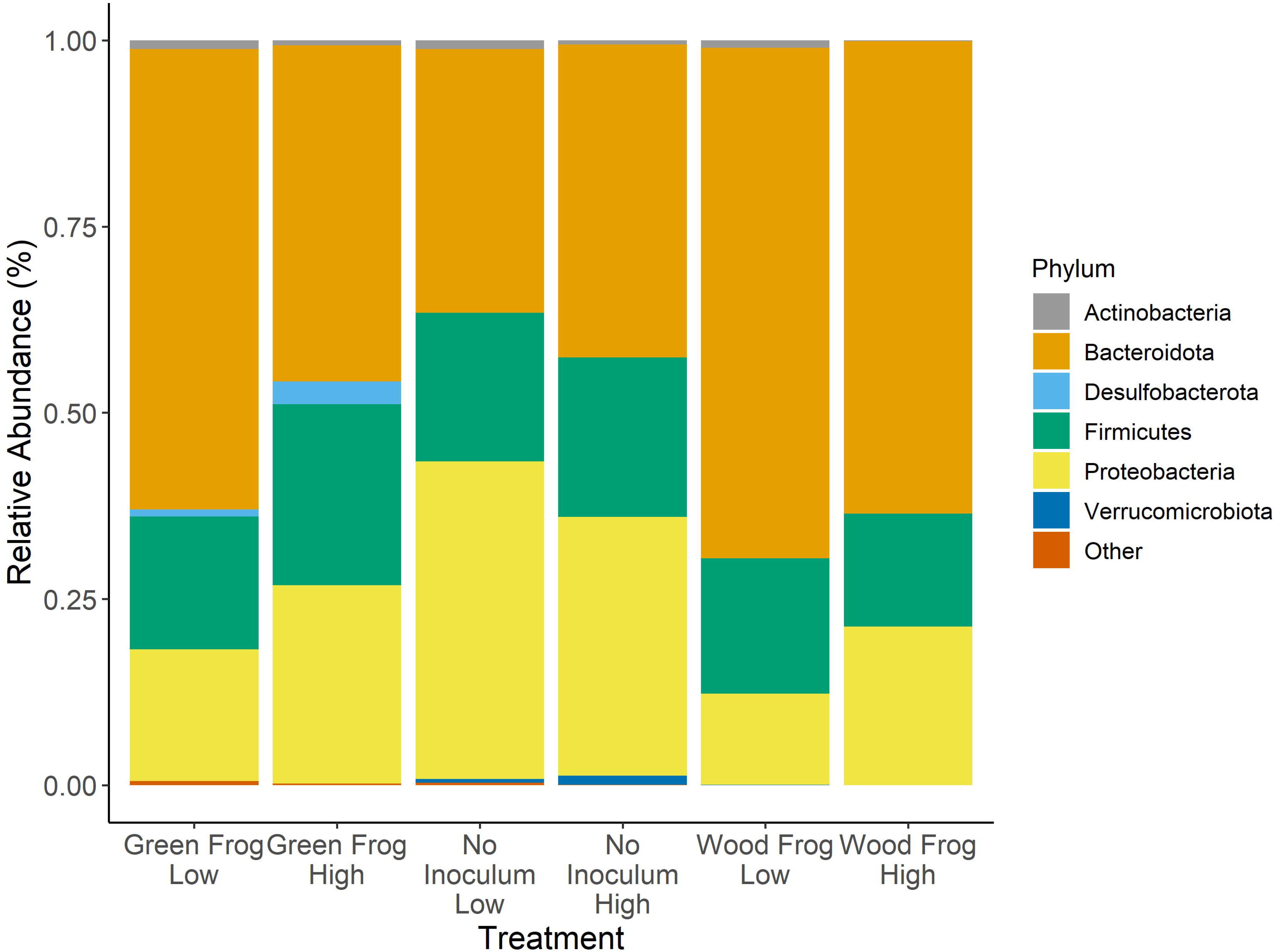
The relative abundance of prominent intestinal bacterial phyla of larval wood frogs that underwent early life microbiota treatments and were acclimated to either low or high temperatures. Larvae that were acclimated to low temperatures are denoted by Low in the X-axis while larvae acclimated to low temperatures are denoted by High.

When examining the relative abundance of bacterial families, only a single family differed between the acclimation temperatures with Beijerinckiaceae being more abundant across all treatment groups at the high temperature (MaAsLin2, corrected P < 0.0001; Table S1). In contrast, there were 14 bacterial families that were significantly different among the three microbiota treatments (MaAsLin2, corrected P < 0.05; Table S1). Specifically, the GF larvae were enriched in Rikenellaceae and *Clostridium* methylpentosum group relative to the other treatments while NI larvae were enriched in Aeromonadaceae.

## Discussion

Increased heat tolerance in organisms is predicted to be beneficial in the future through minimizing the mortality risk to acute heat extremes (Huey et al., 2012). In this study, we build upon current evidence that the gut microbiota influences host heat tolerance. By manipulation of the larval wood frog gut microbiota, we observed that transferring the gut microbiota of a more heat-tolerant larval anuran (green frogs) to wood frogs resulted in elevated heat tolerance, indicating that the microbiota is partially tied to host thermal physiology. These microbiota-manipulated larvae had a higher relative abundance of Rikenellaceae (a SCFA producing bacteria) and a lower relative abundance of Aeromonadaceae (a potential pathogen) when compared to NI larvae. We found a significant, albeit small, effect of short-term acclimation on gut microbiota community composition and diversity suggesting the gut microbiota is susceptible to changes following a brief change in environmental temperature. While our results are correlative, the significance of a cross-species microbiota transplant on increasing the recipient’s CT_max_ underscores the role of the gut microbiota in host thermal tolerance and represents that this method could be useful in species conservation efforts.

Our study is the first to identify that cross-species gut microbiota transplantations influenced the heat tolerance of the recipients in a predictive manner. In line with our prediction, inoculating wood frog eggs with the gut microbiota of larval green frogs, a species with greater heat tolerance than the former (Katzenberger et al., 2021), resulted in higher CT_max_ compared to the WF and NI treatments (Fig. 1). Previous studies in invertebrates have shown that, within species, microbiota transplants of individuals that were acclimated to warmer temperatures promoted enhanced heat tolerance in the recipient (Baldassarre et al., 2022; Doering et al., 2021; Moghadam et al., 2018). Such findings indicate that temperature-based restructuring of the gut microbiota selects for bacterial taxa and/or functions that improve survival under heat stress. For example, Fontaine and Kohl (2023) found that the gut microbiome of larval green frogs exposed to a 24 hr period of heat stress significantly upregulated genes associated with carbohydrate metabolism, transcription, and translation. Surprisingly, they did not find that the heat stress altered bacterial expression of heat shock proteins (HSPs) within the gut microbiota, which are commonly upregulated in response to thermally stressful environments to maintain protein structure integrity (Feder and Hofmann, 1999). This suggests that the gut microbiota improves host heat tolerance through increased metabolic function and DNA processes rather than bacterial HSP expression, but they did not measure intestinal HSP expression, which have been shown to be modulated by the gut microbiota (Arnal and Lalles, 2016). We lack transcriptomic data to explore how the GF treatment promoted greater CT_max_ relative to the other treatments, therefore our conclusions are limited to taxonomic differences we observed rather than altered metabolite and gene expression. Inclusion of metagenomic and transcriptomic analyses of the gut microbiota, along with expression of intestinal HSPs, on studies assessing heat tolerance would be the advisable next step in assessing functional role the gut microbiota has in influencing host CT_max_. Regardless, our findings are in line with other studies showing benefits of cross-species microbiota transplants in species conservation (Dallas and Warne, 2022) and could improve survival for threatened species under warming conditions.

There were several bacterial taxonomic groups that differed among the microbiota treatments. Most prominently, the Bacteroidota family Rikenellaceae, primarily *Mucinivorans hirudinis*, was significantly enriched in GF larvae. This species was initially described in leeches (Nelson et al., 2015), and produces the short chain fatty acid (SCFA) acetate as a byproduct of metabolizing host mucin glycans (Bomar et al., 2011). Its prevalence in other taxa has not received much attention, but *Mucinivorans* species have been previously described in late-stage Asiatic toads (*Bufo gargarizans*) (Chai et al., 2018), suggesting this bacterial species is not limited to leeches. Other members of Rikenellaceae promote beneficial phenotypes such as enhanced heat tolerance in cows (Wang et al., 2022), improved health of late-stage giant spiny frogs (*Paa spinosa*) (Long et al., 2020), and lower obesity rates in humans (Pesoa et al., 2021). One potential mechanism through which *M. hirudinis* could promote heat tolerance is through upregulation of antioxidants as a byproduct of acetate production (Gonzalez-Bosch et al., 2021; Warne and Dallas, 2022). Heat stress is associated with an increased production of reactive oxygen species (ROS) in vertebrates that induce oxidative damage which negatively affect cellular function, host survival, and can limit host CT_max_ (reviewed in Ritchie and Friesen, 2022). Acetate was shown to significantly reduce ROS production in human and mouse cells *in vitro* while promoting greater mitochondrial respiration rates (Hu et al., 2020; Huang et al., 2017). The beneficial role of acetate in promoting antioxidant concentrations has also been demonstrated *in vivo* using piglets (Pang et al., 2021), but direct links between *M. hirudinis* and antioxidant production or ROS reductions are not present. While the link between Rikenellaceae (*M. hirudinis*) and greater heat tolerance is not definitive, its production of acetate could represent a pathway by which GF larvae benefited from its increased relative abundance.

While the WF and NI larvae had similar relative abundances of bacterial families, the latter displayed a higher relative abundance of Aeromonadaceae, specifically those in the *Aeromonas* genus. Members of these family are Gram-negative bacteria that are potential pathogens of amphibians and fish (Janda and Abbott, 2010) and degrade the mucosal barrier of the intestines enabling bacteria to enter systemic circulation of the host (Dong et al., 2018). Additionally, *Aeromonas* can reach relatively high abundances in the intestines of wild (Hird et al., 1983) and lab-reared anuran larvae (Chai et al., 2018). In a study on wild-collected Dybowski’s frog (*Rana dybowskii*), Tong et al. (2020) found that the abundance of *Aeromonas* was enriched in diarrheic adults compared to healthy adults and they posited the presence of this genus lead to the emergence of a pathogenic phenotype. While we did not observe direct evidence of such a phenotype, a reduction in the heat tolerance of NI larvae may be related to higher *Aeromonas* abundance. A potential mechanism for this outcome is through *Aeromonas*-induced systemic-inflammation. As *Aeromonas* metabolites degrade the intestinal barrier (Feng et al., 2022), this could enable Gram-negative bacteria with their lipopolysaccharide (LPS) membrane to enter the bloodstream of the host. LPS promotes inflammation through increased production of pro-inflammatory cytokines and leukocyte activity (Boulange et al., 2016; Kamada et al., 2013; Wang et al., 2020). As an activated immune response can cause a decline in CT_max_ (Hector et al., 2020), the increased Aeromonadaceae abundance in NI larvae may have indicated increased immune costs. This may be indicative that the host could incur negative costs if Aeromonadaceae are enriched in the larval intestines, but further exploration of the topic is required to address this potential relationship.

In addition to increased Aeromonadaceae abundance, the works of Warne et al. (2019) and Fontaine et al. (2022) offer a potential explanation why NI larvae had the lowest CT_max_. These studies demonstrated that larvae experiencing early-life bacterial-depletion had lower mass specific metabolic rates and reduced mitochondrial enzymatic activity, respectively. Fontaine et al. (2022) predicted that larvae with a diminished aerobic scope would have a constrained heat tolerance (Chung and Schulte, 2020; Pörtner et al., 2017), although the importance of such a link has been debated (e.g., Claunch et al., 2021; Ern et al., 2016; Semsar-Kazerouni and Verberk, 2018). Microbial-derived SCFAs are also associated with increased mitochondrial activity (Schonfeld and Wojtczak, 2016), which may constrain the host’s ability to maintain ATP production at high temperatures resulting in a lower CT_max_ (Sokolova, 2023). Compared to GF larvae, NI larvae had a lower, although not statistically significant, relative abundance of Firmicutes and Bacteroidota, two phyla associated with SCFA-production (den Besten et al., 2013; Warne and Dallas, 2022). This could account for our observed CT_max_ pattern. Future studies should examine this link through direct supplementation of SCFAs and then measure the host’s heat tolerance, or by measuring CT_max_ and SCFA concentrations in the intestines. In addition, identifying if changes in the gut microbiota composition affects a host’s aerobic scope would determine if changes in CT_max_ are related to whole-body changes in metabolic rates.

While prolonged acclimation to different thermal regimes alters gut microbiota α- and β-diversity (Baldassarre et al., 2022; Fontaine et al., 2018; Moeller et al., 2020; Theys et al., 2023; Zhu et al., 2021), our short-term acclimation exposure had mixed effects on these metrics. While both bacterial membership and structure were dependent upon acclimation temperature (Fig. 3), only two metrics of α-diversity (Shannon and Simpson indices) displayed a small, temperature-dependent increase (Fig. 2B and 2C). A similar pattern was observed in larval *Ischnura elegans* damselflies exposed to a simulated seven-day heatwave (Theys et al., 2023). The temperature effect in the damselflies was primarily driven by increases in Proteobacteria, a pattern commonly observed in arthropods experiencing warmer temperatures (Sepulveda and Moeller, 2020). In a study on green frog larvae, Fontaine et al. (2022) showed a minor decline in α-diversity with increasing acclimation temperature, but bacterial composition did diverge with the relative abundance of Proteobacteria negatively related to increasing temperatures, although the effect was non-significant. This demonstrates the complex relationship between environmental temperature and the gut microbiota. In line with the predictions of Lozupone et al. (2012) and concept of disturbance ecology (Christian et al., 2015), we propose that exposure to different environmental temperatures represents a disturbance to the gut microbiota that causes a shift in bacterial, diversity, composition, and transcriptomics. While a rapid temperature shift can drastically alter gut microbiota transcriptomics of ectothermic vertebrates (Fontaine and Kohl, 2023; Zhou et al., 2022), community-wide changes are likely more pronounced following prolonged exposure to new temperatures enabling a new community to become established. Longitudinal studies measuring gut microbiota diversity metrics exposed to different acclimation temperatures (Baldassarre et al., 2022) would be beneficial in addressing this prediction.

We found that high temperature-acclimated wood frog larvae were depleted in the Actinobacteria phylum while the Proteobacteria family Beijerinckiaceae (*Methylobacterium-Methylorubrum* sp.) was enriched. The relative abundance Actinobacteria in the gut microbiota, members of which are associated with antibiotic production (Bérdy, 2012), has been shown to be negatively impacted by exposure to warm temperatures in arthropods (Horváthová et al., 2019) and Mongolia racerunners (*Eremias argus*) (Zhang et al., 2022). However, the opposite was found in lab-reared northern leopard frog larvae (*L. pipiens*) (Kohl and Yahn, 2016) and Chinese giant salamanders (*Andrias davidianus*) (Zhu et al., 2021). The pronounced increase in Beijerinckiaceae under the warm temperature was surprising as members of this family have the capacity to grow across a broad thermal breadth in natural environments (Sharp et al., 2014). In previous studies on free-ranging Nile tilapia (*Oreochromis niloticus*) and northern leopard frog larvae, the relative abundance of Beijerinckiaceae (*Methylobacterium* spp.) was enriched at cool temperatures (Bereded et al., 2022; Kohl and Yahn, 2016), while pinfish (*Lagodon rhomboides*) had higher relative abundances at warmer temperatures (Givens, 2012). Many Beijerinckiaceae members are associated with nitrogen fixing (Marin and Arahal, 2014) which can provide their host with additional energy (Russell et al., 2009), although how this influences larval anuran performance or physiology requires further assessment. Our results show that complex relationships exist between environmental temperature and the relative abundance of Actinobacteria and Beijerinckiaceae that are likely host specific.

A limitation of our study design is that we did not have an unmanipulated microbiota treatment that represents wildtype wood frog larvae. The WF treatment was meant to restore the wildtype microbiota following antibiotic exposure, but a subsequent study showed that unmanipulated wildtype larvae significantly diverged in both alpha and beta diversity indices from WF larvae (Dallas et al., In Prep.). Therefore, the WF larvae did not accurately represent the wildtype condition. While GF larvae exhibited the highest CT_max_ among our microbiota treatments, their CT_max_ were comparable to less-developed wildtype larvae (Fig. S1). This indicates that transplanting the green frog microbiota improved heat tolerance of recipient larvae relative to the other antibiotic-treated larvae but not unmanipulated individuals. As early-life antibiotic exposure in anurans leads to phenotypic costs including higher mortality risk, delayed metamorphosis, and increased tail deformities (Peltzer et al., 2017; Warne et al., 2019; Zhu et al., 2022), suggesting that the green frog transplantation rescued antibiotic-treated larvae from the lower heat tolerance evident in the NI and WF larvae. A follow-up study incorporating a more successful within-species microbiota inoculation would provide greater support for linking cross-species gut microbiota transplants to warmer CT_max_.

## Conclusion

As the risk of extreme heat events becomes more prevalent under climate change, how the gut microbiota modulates host thermal physiology represents an intriguing research question. Our results represent an important step in linking this relationship by demonstrating that inoculating antibiotic-treated wood frog larvae, a heat-sensitive species, with the gut microbiota of green frog larvae, a more heat-tolerant species, promoted higher CT_max_ relative to other antibiotic-treated larvae. This suggests that interspecific differences in heat tolerance are partially driven by the gut microbiota, and that this difference can be transferred between species simply through microbial transplants. The green frog transplant resulted in large increases in the Bacteroidota family Rikenellaceae, a producer of the SCFA acetate, which is linked to increased antioxidant activity (Gonzalez-Bosch et al., 2021). As heat shock increases the production of ROS that disrupt cellular and mitochondrial activities which restrict heat tolerance, a greater abundance of antioxidants can maintain cell function at high temperatures. However, we acknowledge that our results are correlative but the inclusion of metagenomics to assess a mechanistic link between the gut microbiota and heat tolerance represents a future direction to explore. Microbial transplants represent a species-rescue technique against multiple threats, and this study represents that cross-species transplants can rapidly induce greater heat tolerance which promotes survival under warming temperatures.

## Acknowledgments

We would like to thank Jacob Hutton for his assistance in collecting the larval green frogs used in the microbiome treatment, and both Jacob and Adrian Macedo for their assistance in the CT_max_ assays. We also wish to acknowledge Dr. Frank Anderson and Dr. Derek Fisher for their comments on previous versions of this manuscript.

## Data Availability

Data is available upon request.

## Table and Figure Legends

**Figure S1:** The critical thermal maximum (CT_max_) of larval wood frogs across all gut microbiota treatments and unmanipulated wildtype larvae. Center lines within boxplots represent the median and the boxes denote the interquartile range with whiskers representing 1.5× the upper or lower quartile.

**Table S1:** Relative abundances of bacterial phyla and families in tadpole gut microbial communities that were significantly impacted by acclimation temperature and gut microbiota treatment. Relative abundances are displayed as means ± 1 standard error. The group in which the specific taxa was most abundant is in bold. Statistical testing was conducted using MaAsLin2. N.O. represents groups where the bacterial taxa was not observed. P-values were corrected using the BH FDR method.

